# Rapid genome-wide introgression reveals fitness advantage of immigrant genotypes

**DOI:** 10.1101/2025.08.27.672692

**Authors:** Ben A. Flanagan, Arshad Padhiar, Foen Peng, Saif Quraishi, Andrea J. Roth-Monzón, Fahad Gilani, Lauren Simonse, Meghan F. Maciejewski, Noah Reid, Milan Malinsky, Amanda K. Hund, Daniel I. Bolnick

**Affiliations:** Department of Ecology and Evolutionary Biology, University of Connecticut; 75 North Eagleville Rd, Storrs CT 06269 USA; Current affiliation: Department of Biology, Haverford College; 370 Lancaster Ave, Haverford PA 19041 USA; Current affiliation: Chicago College of Osteopathic Medicine; 555 31^st^St., Downers Grove IL 60515, USA; Current affiliation: Museum of Vertebrate Zoology, University of California Berkeley; 3040 Valley Life Sciences, Berkeley CA 94720-3140 USA; Current affiliation: School of Integrative Biology, University of Illinois, 505 S. Goodwin Ave, Urbana, IL 61801, USA; Department of Molecular and Cell Biology, University of Connecticut; 75 North Eagleville Rd, Storrs CT 06269 USA; Institute of Ecology and Evolution, University of Bern; Bern, Switzerland; Department of Biology, Carleton College, Northfield, Minnesota, USA

## Abstract

Evolutionary biology has long recognized the tendency for populations to be locally adapted to their ancestral habitat, resulting in higher resident fitness. However, immigrants can also introduce beneficial alleles. The resulting adaptive introgression is usually inferred retrospectively, rather than as a contemporary process. Here, we document exceptionally rapid ongoing adaptive introgression in a lake population of threespine stickleback (*Gasterosteus aculeatus*). In the first generations after a discrete immigration event, all chromosomes exhibited large increases in immigrant ancestry due to linkage disequilibrium. After a decade, the extent of introgression varied across the genome. The fastest-evolving genes included *Spi1b*, which enables an increased fibrosis defense against a previously common tapeworm, whose prevalence then declined dramatically. This case study highlights the capacity for immigration to supply beneficial alleles that drive rapid genome-wide evolution.

## Main Text

Evolutionary theory leads us to expect local adaptation – the tendency for natives in a particular environment to have higher fitness than immigrants (*1*). Generations of natural selection retain locally beneficial alleles and remove locally harmful alleles, whereas immigrants carry alleles favored in their native range but untested in their new environment. The resulting selection against immigrant genotypes (*2*) facilitates population divergence and speciation (*3*). Yet, immigrants can outperform residents (*4*). They may import universally beneficial alleles that arose elsewhere in the species range, or may carry alleles pre-adapted to changing local environments. For instance, if parasites gain an edge during coevolution, then resident hosts can be locally maladapted compared to immigrants (*5*). Once introduced, beneficial immigrant alleles can increase in frequency leading to adaptive introgression (*6*). Examples of adaptive introgression exist in plant and animal species (*7–9*), and often entail introgression of alleles at immune genes such as MHC, conferring enhanced resistance to locally adapted parasites (*10*– *12*). However, we know little about the earliest phases of adaptive introgression, which is usually inferred long after the actual event by identifying genomic regions with atypical ancestry (e.g., Neanderthal sequences in modern human genomes (*13*)). This retrospective inference makes it difficult to determine the exact timing and speed of introgression, and its initial genomic footprint. Here, we use genome sequences spanning two decades to document a contemporary example of adaptive introgression between conspecific populations of threespine stickleback. We reveal very strong selection in the earliest generations of introgression, driving genome-wide increases in immigrant ancestry for all chromosomes. In later generations, introgression began to differ among chromosomes and loci allowing us to identify likely targets of selection, such as a gene conferring immunity to a formerly abundant tapeworm.

The threespine stickleback (*Gasterosteus aculeatus*) is a small fish found in northern coastal habitats. After Pleistocene deglaciation, marine stickleback repeatedly invaded and then adapted to freshwater habitats, including loss of armor plating, and gain of resistance to freshwater parasites (*14*, *15*). As the many isolated lake populations adapted to their respective habitats and communities, they diverged in morphology (*16*), and immunity (*17*). For some traits this among-population divergence evolved in parallel, creating replicated trait-environment correlations that strongly implicate a role for natural selection (*18*). Furthermore, effective population sizes within lakes are large, so genetic drift should be relatively weak (*19*). Yet, evidence for local adaptation remains mixed. One reciprocal transplant experiment moved fish between cages in lake and stream habitats, finding higher resident fitness within one watershed, but not other watersheds (*20*). A second transplant experiment found that one population outperformed another in all environments (*21*). A third found evidence for local maladaptation: stickleback confined to cages in their native habitat (eg., lake fish in their lake, stream fish in their stream) were more heavily infected by parasites than immigrants (*22*). These results suggest that local maladaptation can be common (and asymmetric), potentially favoring adaptive introgression. Theory suggests that parasitism and immune evolution may be an especially potent force for such introgression (*5*), and stickleback have evolved substantial population differences in immune gene alleles (*17*), expression (*23*), and immune responses (*24*).

Here, we report a genomic time-series revealing exceptionally rapid adaptive introgression in a stickleback population from Gosling Lake, Vancouver Island. A 2005 survey of lake stickleback across Vancouver Island identified Gosling Lake as having unusually high prevalence of a tapeworm, *Schistocephalus solidus* (*25*). Subsequent genetic mapping revealed that Gosling Lake (GOS) stickleback had evolved infection tolerance: selection had driven fixation of a 78-bp deletion which eliminated a predicted CTCF binding site in the gene *spi1b* (*24*). This *spi1b^del^* allele was associated with a tolerance strategy: GOS fish fail to develop fibrosis after infection, whereas the ancestral *spi1b^+^* was associated with a fibrosis response that suppressed tapeworm growth. After post-glacial colonization of Gosling Lake, selection had favored *spi1b^del^* over the fibrosis-prone ancestral *spi1b^+^* allele, apparently reflecting a tolerance strategy (*24*). Although GOS were fixed for *spi1b^del^* in 2009, we were surprised to find *spi1b^+^* homozygotes in a 2018 sample of GOS fish caught to generate embryos. PCR reactions to detect the deletion confirmed that *spi1b^+^* had increased in frequency between 2009 and 2018. We therefore tested for evolution genome-wide, by comparing sequences of 2009 versus 2022 samples from the lake (N=100 and 108 respectively). This comparison revealed large shifts in allele frequencies on all chromosomes (mean between-year F_ST_=0.175, mean allele frequency change Δp=0.227, Fig. S1), in 13 generations (or fewer). At many loci, these changes involved alleles novel to the GOS fish (absent in the 2009 sample), suggesting an effect of introgression rather than selection on standing variation.

To identify possible sources of introgression, we compared allele frequencies from the 2022 GOS fish genomes, with allele frequencies from 36 populations throughout Vancouver Island from a previous ddRAD study (*18*). Previously, lakes within the Campbell River watershed (including Gosling) had formed a well-supported clade, distinct from neighboring watersheds. Yet, in 2022 GOS fish were as closely related to lakes in the isolated Mohun watershed (F_ST_=0.125 between GOS and Comida Lake [COM] 9.5 km away), as they were to their immediate neighbor Boot Lake (1.3 km away, with F_ST_=0.123; Fig. S2 & S3). The recent GOS genotypes bore less affinity to other lakes in the region, suggesting introgression from the Mohun watershed (Comida and Mohun Lakes, COM and MOH). In a 2010 experiment testing for local adaptation, stickleback from COM and MOH had been transplanted into cages in Gosling Lake (*23*) (with provincial approval Fish Transfer Permit IT-12085). Some cages in Gosling Lake were vandalized, inadvertently releasing 37 COM and 15 MOH fish into Gosling Lake. This is a large lake (62.5 ha, 6.6 km of shoreline), so the effective population size of the residents is in the tens of thousands (*19*) and the census size of adults is likely in the hundreds of thousands or more (a survey of nests found >1 nest per meter of surveyed shoreline (*26*)). Therefore, the inadvertently introduced fish constituted less than a tenth of a percent of the resident population.

Re-sequencing additional individuals from 2007-2022 GOS collections (and five nearby populations; Table S1) confirmed that the Mohun watershed (COM and MOH) was the likely source of immigration (Fig. 1A). The timing and speed of introgression imply exceptionally strong selection, with Mohun watershed ancestry increasing dramatically between 2010 and 2013 and continued to increase thereafter (Fig. 1B). From 2005 to 2010 all sequenced fish exhibit pure GOS genotypes. In 2011 and again in 2012 we sampled one individual GOS-Mohun hybrid (Fig 1C; Fig. S4), the remaining 29 fish per year were 100% GOS (Fig. 1D). One generation later (2013) GOS-Mohun heterozygotes outnumbered native GOS homozygous genotypes on all chromosomes (Fig. 1E). By 2022 most chromosomes exhibited a majority of introgressed Mohun watershed homozygotes, though the extent of introgression varied among chromosomes (Fig. 1F). The speed of the introgression can be seen by plotting the proportion immigrant ancestry through time for each chromosome (Fig. 1G). Through 2018, there were no significant between-chromosome differences in the proportion immigrant ancestry (P>0.05), confirming that the speed of early introgression was similar genome-wide.

**Figure 1:**
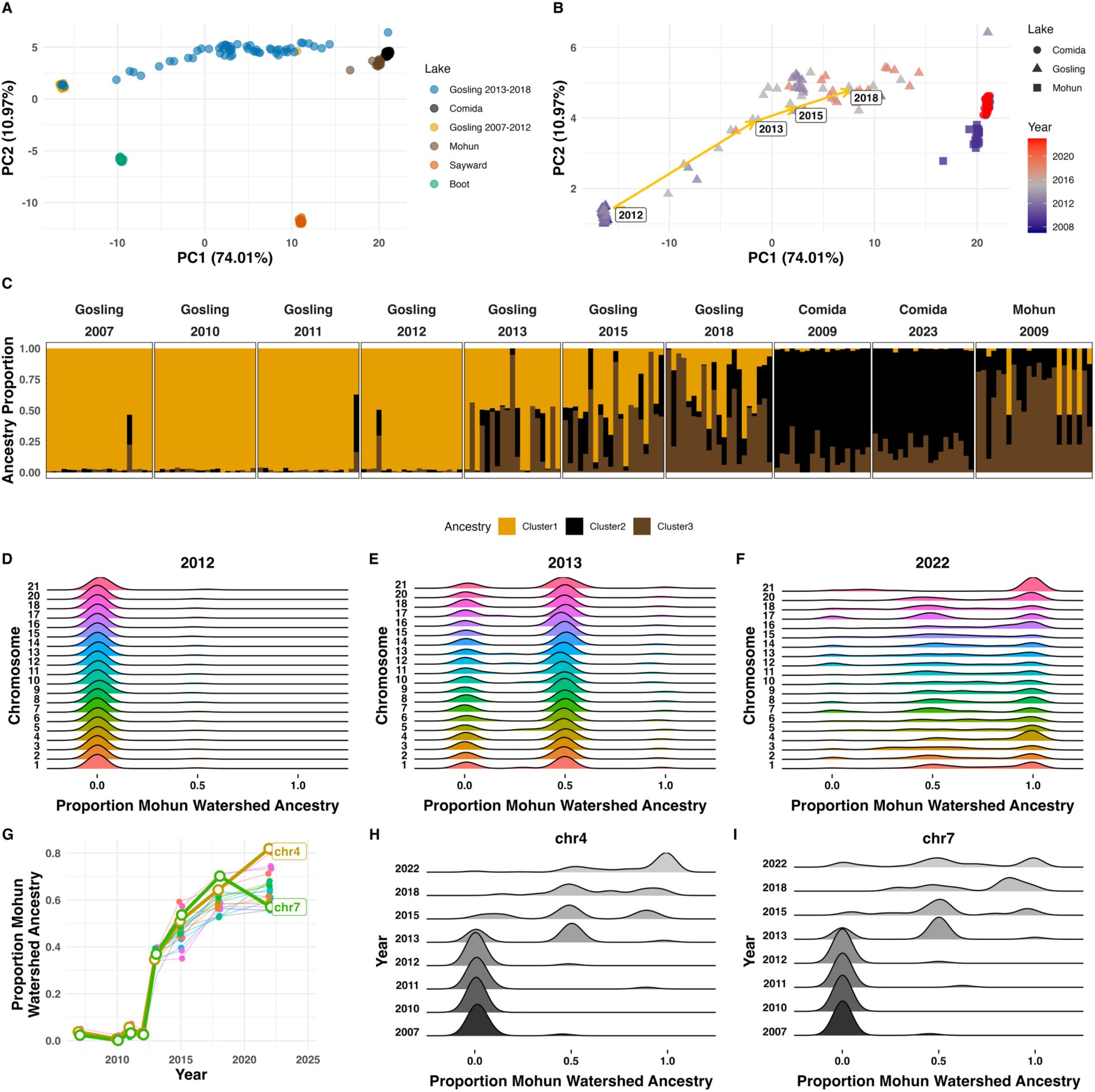
Rapid genome-wide introgression. (**A**) Genome-wide principal components analysis for Gosling (GOS) samples from 2007-2018 (lcWGS sequence data), along with Comida Lake (COM), Mohun Lake (MOH) and a sample of nearby lakes and marine outgroup (Sayward). (**B**) Time course of introgression with Gosling and Mohun watershed lakes. Labels indicate annual averages. (**C**) Admixture estimations for Gosling Lake, and the Mohun Watershed (COM, MOH) for k = 3. (**D - F**) The Mohun watershed ancestry proportion in Gosling Lake samples for each chromosome in the years 2012, 2013, 2022. (**F**) Mean proportion of Mohun watershed ancestry over time, for each chromosome. (**G-H**) Time course of Mohun watershed ancestry proportion for Chr4 and Chr7.

Normally studies of adaptive introgression reveal genomic windows of introgression: sites subject selection that drives a beneficial new allele (and closely linked sites) into the recipient population; this is a mirror image of the ‘genomic islands’ of divergence observed during population differentiation (*27*). However, in the earliest generations of introgression there is little time for recombination between physically linked sites or even for independent assortment of chromosomes. Consequently, for the first generations the entire genome behaves as if it were linked, a phenomenon not observable in retrospective studies of adaptive introgression’s long-term effects. Specifically, if selection of strength *s* favors an immigrant allele at one locus, even neutral loci on other chromosomes experience hitchhiking selection of strength *s/2^g^* in generation *g*. The spread of Mohun watershed ancestry in the GOS population fits this model closely, with an effective selection coefficient *s*=0.29 (Fig. S5). This is exceptionally strong natural selection, favoring immigrant genotypes at one (or more) loci.

In later generations, independent assortment erodes disequilibrium between chromosomes, and the effective strength of selection on neutral loci *s/2^g^*asymptotically approaches zero (*28*). Consistent with this theory, after ∼10 generations, many chromosomes leveled off at 64% Mohun watershed ancestry. However, selection should continue to drive introgression on chromosomes carrying the beneficial immigrant variants. Conversely, any loci that originally harbored locally adapted alleles might cause their chromosomes to reverse direction back towards greater resident ancestry. We indeed see that, by 2022, the extent of introgression differed among chromosomes (Fig. 1G, Type II ANOVA, chromosome*year F_20, 5294_=4.387, P <0.001; Figs. S6&S7). Chr4 approached 80% Mohun watershed ancestry (Fig. 1H). In contrast, Chr7 reversed and became more GOS-like between 2018 and 2022 (Fig. 1I), suggesting this chromosome harbored locally adapted native alleles, that had been transiently overwhelmed by linked selection for immigrant genotypes elsewhere.

This emerging among-chromosome variation in introgression helps us identify chromosomal regions, and perhaps candidate genes, most likely to be targeted by selection for immigrant alleles. Locus-specific F_ST_ (2009 versus 2022) varied substantially across the genome ranging from 0.0 to 1.0 (Fig. 2A; median F_ST_=0.1155, mean F_ST_=0.1754, st.dev.=0.176, 90% quantile = 0.4561, 95% quantile = 0.5732, 99% quantile = 0.963, n = 1884931). Loci with especially high F_ST_ (e.g., above the 95% quantile) represent candidate targets of selection which includes many variable sites (n = 94247). The genes closest to the top 5% F_ST_ loci were enriched for molecular function ontologies related to small molecule binding (GO:0036094), catalytic activity (GO:0003824), transporter activity (GO:0005215), and trace-amine associated receptor (TAAR) activity (GO:0001594). Some of the most-divergent loci are the TAAR genes (Fig. 2A), 13 of which occur on Chr16, which shows exceptional genetic change over 10 generations (Fig. 2B). The TAAR gene family plays a role in immune function, being required for innate cells called granulocytes to chemotactically migrate towards parasites (*29*). TAARs function in the vertebrate gut (*30*) which is the site of *S. solidus* invasion in stickleback, so this gene family may play an important role in resistance during early stages of stickleback-tapeworm interactions.

**Figure 2:**
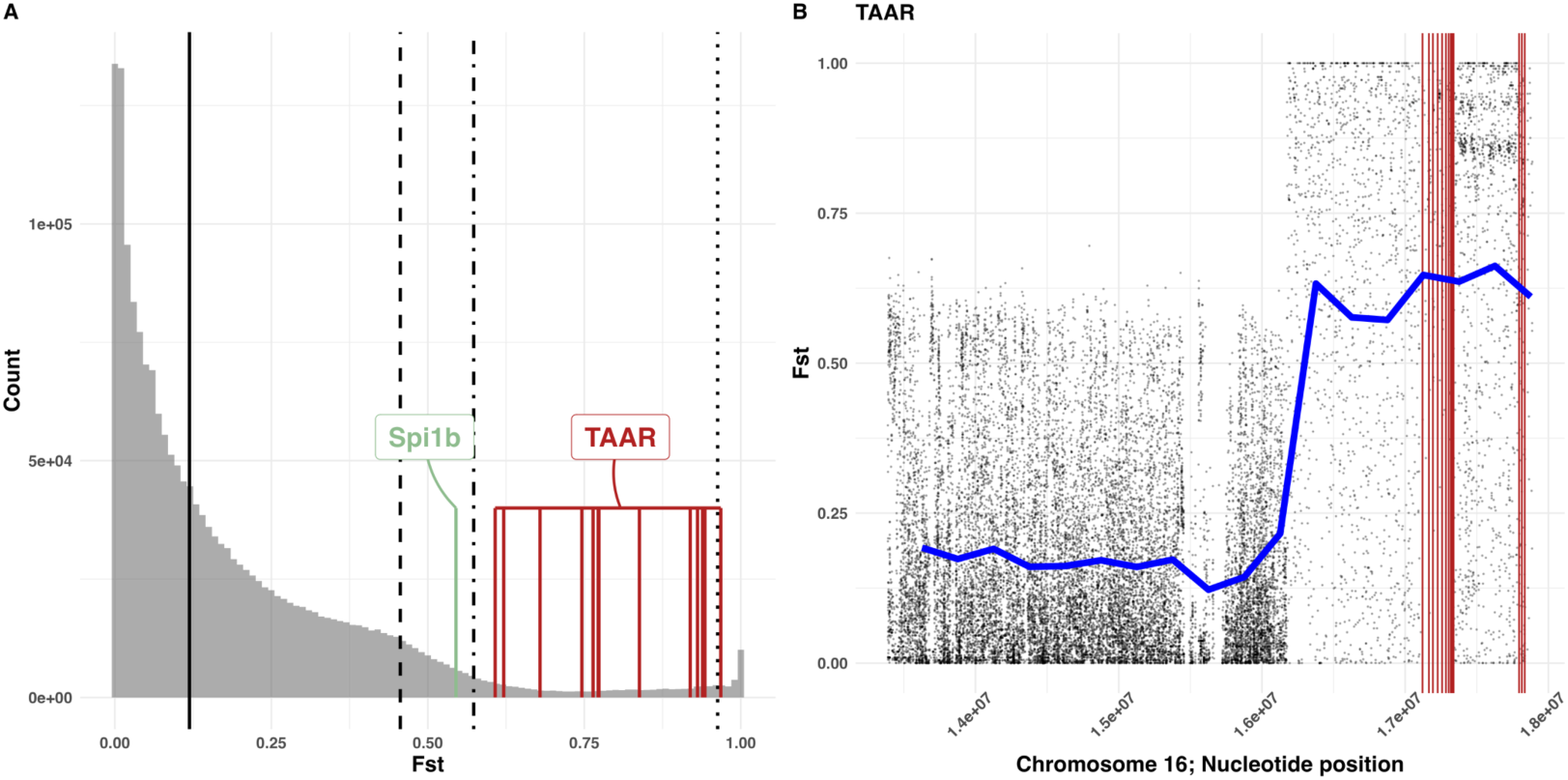
Genetic differentiation spanning the introgression event in Gosling Lake from 2009 versus 2022. (**A**) Histogram of between-year FST values. The vertical solid line is the genome-wide mean FST, the dashed line indicates 90th quantile, the dash-dotted line represents the 95th quantile, and the dotted line represents the 99th quantile. The red lines indicate FST values for SNPs closest to trace-amine associated receptor (TAAR) genes (GO:0001594), the pale green line indicates FST for *Spi1b*. (**B**) Fst surrounding the TAAR loci (red) on Chr16 with smoothed FST trends in blue.

Previous genetic mapping suggested that the gene *Spi1b* influences the stickleback fibrosis response to tapeworm infection (*24*). The *Spi1b^del^* allele which was fixed in GOS fish, was genetically correlated with a suppressed fibrosis response, probably an example of evolved tolerance. Between 2005-2010, all 320 sampled GOS fish were *Spi1b^del^*homozygotes (Fig. 3A, allele frequency=1.0 [0.9882, 1.0]). But, *Spi1b* was among the top 10% fastest-diverging genes after 2010 (Fig. 2A, F_ST_=0.545). Using PCR genotyping, we found the first *Spi1^+^/Spi1b^del^*heterozygote in 2012 (frequency of 1.5%). In just a single year after this (one generation), the *Spi1b^+^* allele frequency increased to 40% in 2013. In a decade the immigrant allele reached a frequency of 58.3% by 2022 (Fig. 3B; Table S2). In 2022, individuals with the *Spi1b^+^* allele had disproportionately more COM ancestry (Fig. 3C). Although Chromosome 2 as a whole did not show exceptional introgression (Fig. 1F), local ancestry PCA (*31*) reveals especially high COM ancestry in the narrow chromosomal region immediately surrounding *Spi1b*, relative to the rest of Chr 2 (Fig. 3D). From this, we infer that *Spi1b* is likely to be one of the targets of selection during introgression, with the immigrant *Spi1b^+^* allele partially replacing the *Spi1b^del^* allele that had been inferred to provide fibrosis suppression and tolerance of *S. solidus* infection (*24*).

**Figure 3:**
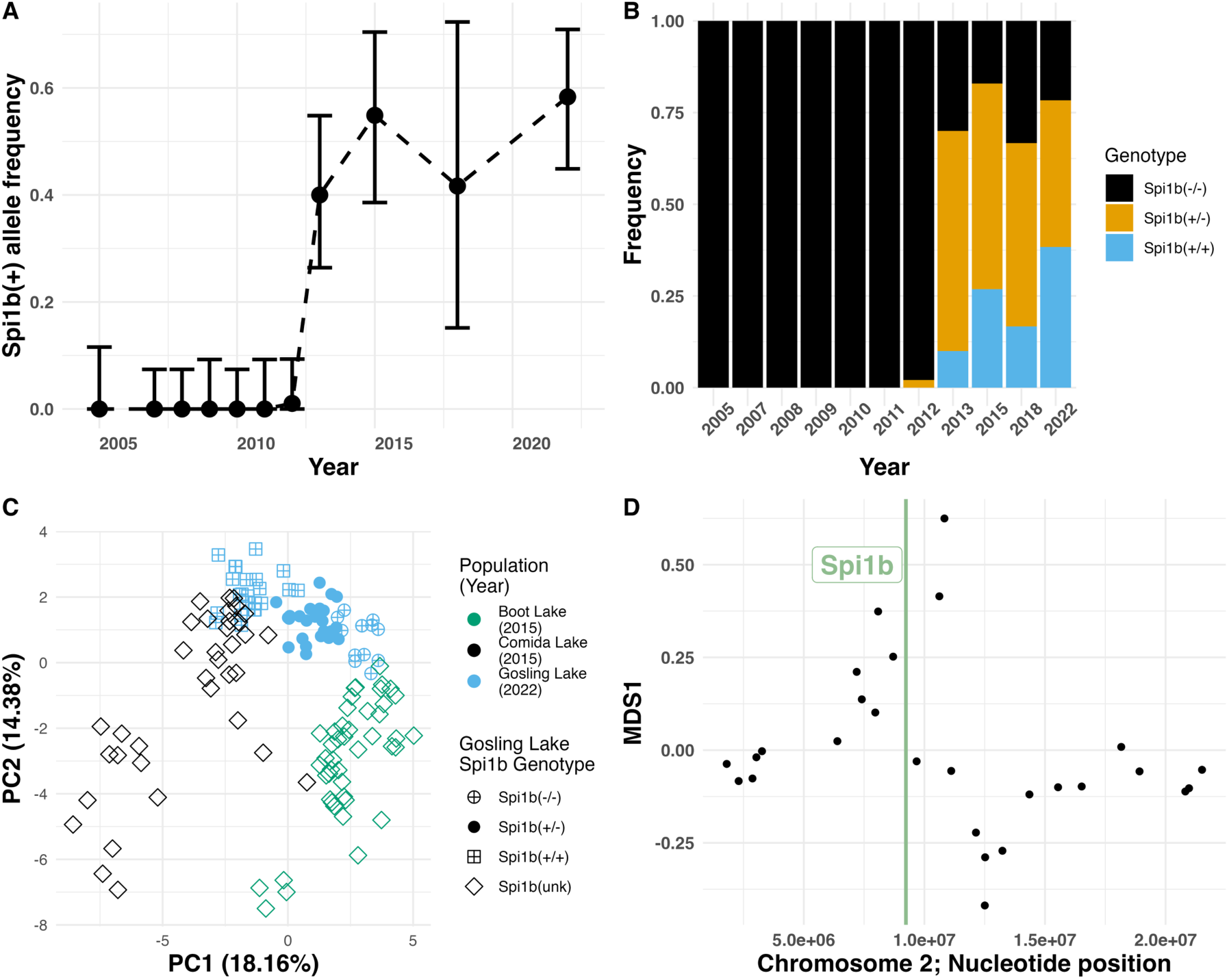
Introgression of a candidate gene, *Spi1b*. (**A**) Spi1b^+^ allele frequency change during the pulse of introgression, (**B**) genotype frequency change prior and following the introgression event, (**C**) association of *Spi1b* PCR based genotype for Gosling Lake with ancestry PCA. (**D**) the variation in structure between windows across Chr2 for Gosling Lake sampled in 2022 with Comida Lake, and Boot Lake. Each point represents a non-overlapping genomic window (mean window size 539.3kb with three loci) wherein a PCA is performed uniquely for each window then window similarity is scored using multidimensional scaling (MDS) transformation. The genomic windows adjacent to Spi1b (vertical red line) show dissimilar population structure in MDS space relative to the background structure present on Chr2.

The introgression of Mohun watershed ancestry (including *Spi1b*) coincided with a steady decline in tapeworm prevalence in GOS stickleback. Tapeworm prevalence in GOS dropped from 73% (2005) to just 8% (2022) (Fig. 4A; binomial GLM, logRR=–0.18, P <0.0001), negatively associated with increased *Spi1b^+^* allele frequency (Fig. 4B). Meanwhile, fibrosis increased transiently in GOS fish (Fig. 4C). Early surveys of Gosling Lake stickleback (2005, 2009, 2010, 2011) detected no fibrosis despite the high prevalence of infection. This matches previous reports that lab-raised GOS fish lacking the fibrosis response to *S. solidus* (*24*).

**Figure 4.**
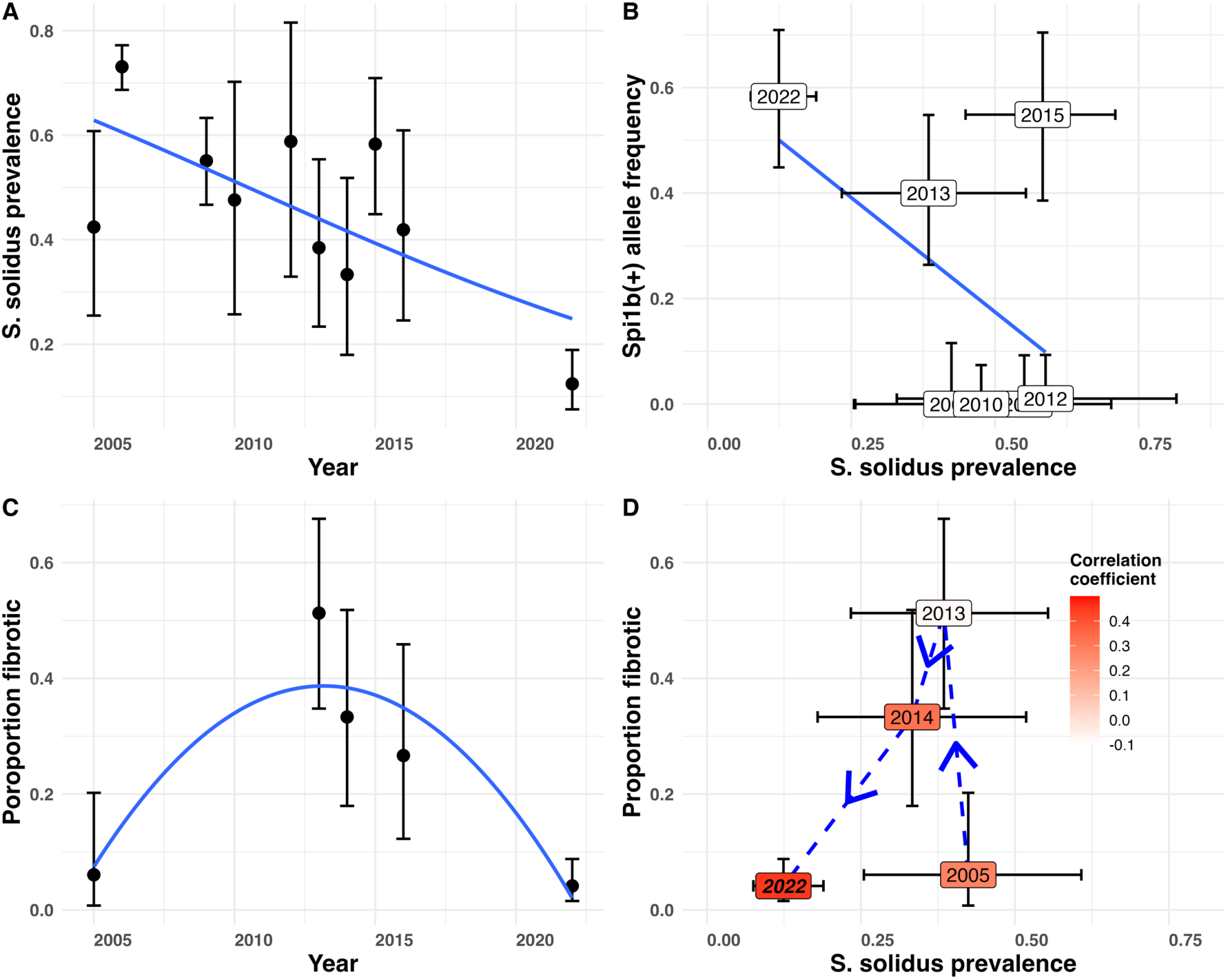
Temporal dynamics of parasite prevalence, *spi1b* with a 78-bp deletion allele frequency, and the fibrosis frequency in Gosling Lake. (**A**) From 2005 to 2022 the *S. solidus* prevalence decreased over time during the introgression. (**B**) Changes in *spi1b* reference allele frequency were associated with changes in tapeworm prevalence during the introgression period. (**C**) Peritoneal fibrosis exhibits a significant quadratic change in prevalence over time, from initially low fibrosis, to high fibrosis early in introgression, to low fibrosis when the parasite is less common. (**D**) A trajectory of the relationship between parasite prevalence and fibrosis frequency over time, informed by the immune eco-evo dynamic model presented in (*32*). The shading within each time point illustrates the strength of correlation between log-transformed *S. solidus* prevalence and fibrosis score. The bold italics font face in 2022 signifies a significant correlation (P <0.05).

However, in 2016 we were (at the time) surprised to observe severe fibrosis in 9 out of 31 sampled fish. By 2022, fibrosis was present and disproportionately found in the few fish with infections (correlation between log infection intensity and fibrosis score, r=0.449, df=143, P<0.0001) (Fig. 4D). These results confirm that GOS stickleback gained the capacity for fibrosis response, which had been absent a decade earlier (*24*). Plotting the trajectory of infection and fibrosis over time (Fig. 4D), we see a shift from high infection and low fibrosis (2005) to high fibrosis (2013), then reduced infection and reduced parasitism in 2014, and even less in 2022.

This counter-clockwise arc in Fig. 4D matches theoretical expectations for the evolution of an inducible immune response (*32*): an initially susceptible population is heavily infected, but as inducible immunity evolves, infections decline. At low infection rates, the hosts retain the genetic capacity for an immune response but are no longer triggered to express the trait. As parasite abundance declines, selection for the immune trait weakens so the defensive allele stops increasing in frequency (*32*). This model thus can explain why selection for *Spi1b^+^*, though initially strong, may have weakened in the later generations.

Most studies that use outlier scans to detect targets of selection in wild populations typically do not confirm candidate genes’ function (*33*). However, one of our putative targets of adaptive introgression, *Spi1b*, has previously been linked to fibrosis and tapeworm resistance during studies of this same population (*24*). Two new experiments confirm that previous QTL association. First, we performed CRISPR/Cas9 gene editing to knock out *Spi1b* in F1 hybrid embryos from a Gosling × Roberts cross generated in 2018. All surviving F2 progeny of these F1s (n>300) were heterozygous knockouts (*Spi1b^+^/Spi1b^KO^*), so *Spi1b* knockouts are lethal when homozygous. When the edited fish were a year old, we used alum injections to induce fibrosis (*34*) in heterozygous knockout fish and wild type siblings, with saline control injections. At both 7 and 28 days post-injection (DPI) *Spi1b^+^/Spi1b^KO^* knockout fish exhibited significantly elevated fibrosis compared to *Spi1b^+^/Spi1b^+^*siblings (Fig. 5A&B; Alum F₁,₇₀=350, P <0.001; DPI F₁,₇₀=343, P <0.001; knockout F₁,₇₀=5.7, P=0.020; Alum×DPI F₁,₇₀=250, P <0.001). This significant effect of *Spi1b* editing confirms that this gene contributes to fibrosis, but the effect direction was opposite to our initial expectation (the 78-bp deletion having been associated with lower fibrosis). This unexpected effect direction may be due to the use of a knockout that does not recreate the 78bp deletion hypothesized to effect gene regulation. Additionally, the immune adjuvant (alum) used to induce fibrosis evokes somewhat different transcriptional pathways when compared tapeworm infection or tapeworm protein injection (*35*).

**Figure 5:**
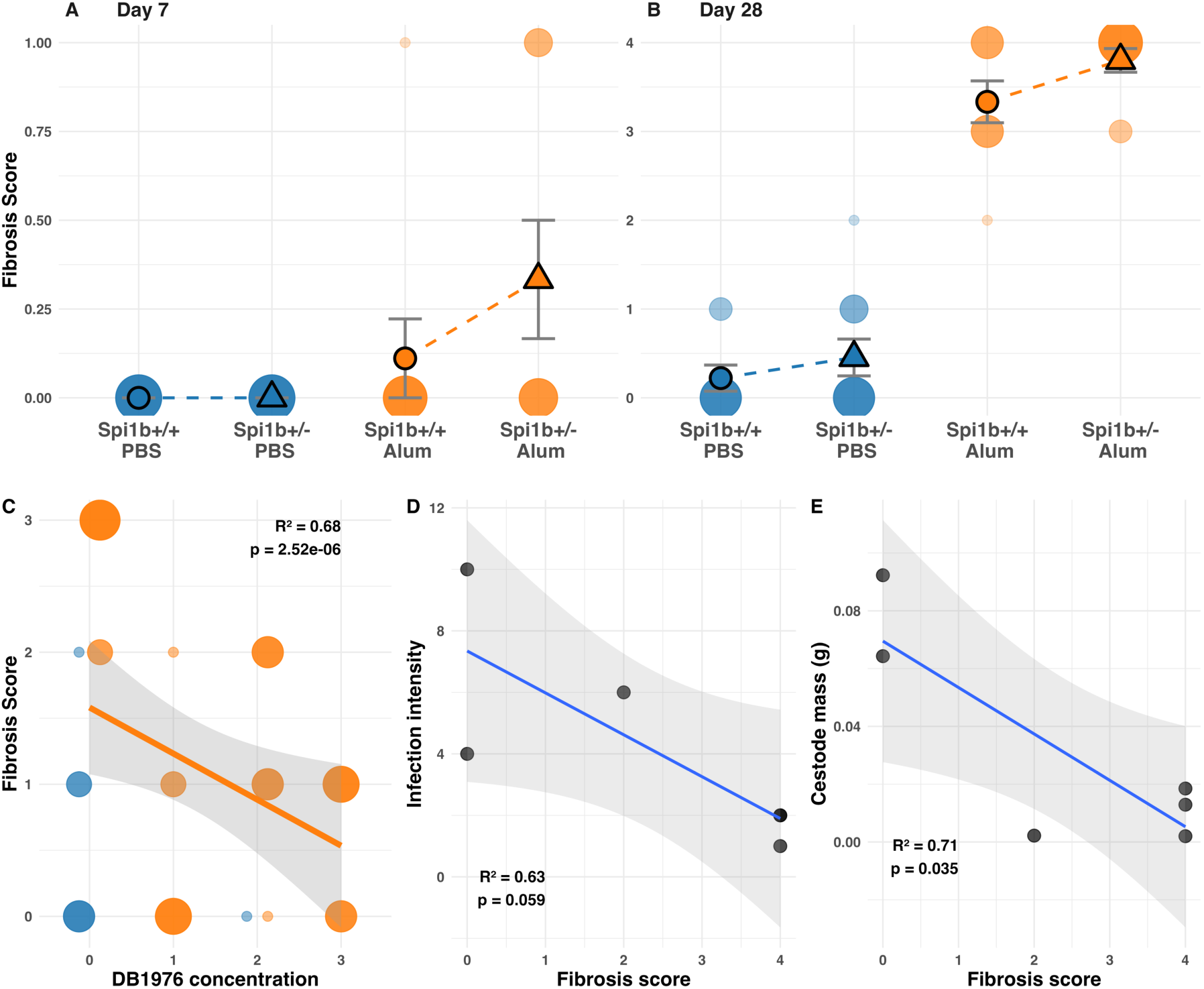
Spi1B expression regulates the fibrosis immune response in Stickleback. The association between *Spi1b* knockouts and fibrosis in response to alum or saline injections (PBS) at (**A**) 7 and (**B**) 28 days post-injection. (**C**) db1976 inhibition of *Spi1b* causes suppression of fibrosis in marine stickleback injected with alum. This inhibition of fibrosis is dose-dependent (n=8 for each concentration). Alum-induced fibrosis suppresses tapeworm infection (**D**) intensity and (**E**) mass in lab-raised GOS stickleback experimentally fed 10 tapeworms.

Pharmacological inhibition of *Spi1b*’s protein product further confirms our conclusion that *Spi1b* plays a key role in fibrosis and better fits our expected effect direction. *Spi1b* produces a protein called PU.1, a transcription factor activating fibroblasts (*36*). The small-molecule inhibitor db1976 blocks PU.1 activity (*37*), avoiding potential artifacts of gene editing. The pro-inflammatory adjuvant alum induces fibrosis in all stickleback populations regardless of genotype (*34*) (t=4.94, P <0.0001), and nearly all fish species (*36*). We co-injected a marine population of stickleback with alum (or saline controls) with varying doses of db1976 (or DMSO controls). Importantly, db1976 reduces the fibrosis response to alum in a dose-dependent manner (Fig. 5C; t=–3.83, P=0.0007). These results confirm that *Spi1b* affects sticklebacks’ peritoneal fibrosis response.

We then experimentally confirmed that fibrosis plays a causal role in tapeworm resistance. Immediately after alum injections to induce fibrosis in lab-raised Gosling Lake fish (lab raised from 2019 wild caught parents, genotype unknown) we fed individuals tapeworm-infected copepods. The pharmacologically-exaggerated fibrosis reduced both the number and mass of tapeworms compared to PBS-injected controls (Fig. 5D; Poisson GLM, Z=–2.78, P=0.005 for infection intensity; Fig. 5E; r=–0.71, P=0.035 for tapeworm mass).

Our observations document remarkably rapid genome-wide adaptive introgression caught in real time through long term genomic sampling. A small number of immigrants contributed foreign alleles into a large established population, likely in 2010. In less than a decade, the immigrant genotypes had increased to become the majority. Contrary to the common expectation in evolutionary biology that immigrants between conspecific populations will be locally maladapted in their new environment, this rapid introgression clearly shows that gene flow can supply beneficial genetic variants favored by strong natural selection. Such strong selection drives genome-wide introgression in the earliest generations, affecting all chromosomes essentially equally. This genome-wide effect is missed in retrospective studies inferring ancient admixture, which left only scattered fingerprints of Neanderthal ancestry (*13*). This is because over later generations, selection continued to drive introgression at some loci and chromosomes, but other chromosomes leveled off or even began to revert towards native genotypes. We infer that strong initial selection is followed by a sorting process once independent assortment and recombination sufficiently reduce linkage disequilibrium so that selection can act separately on immigrant and locally adapted loci. The targets of continued introgression included loci linked to parasite resistance. In particular, the fibroblast-activating gene *Spi1b* was previously associated with stickleback resistance to *S. solidus* tapeworms through genetic mapping (*24*) and transcriptomics (*35*). The introgression of *Spi1b*^+^ was faster than much of the rest of the genome, especially early on, and we see a corresponding increase in fibrosis and decline in *S. solidus* prevalence. We experimentally confirmed the effect of this gene through cas9 knockouts, drug inhibition, and an infection experiment. From these results we infer that part of the introgression (though by no means all) was likely driven by increased immunity to the abundant and highly virulent *S. solidus* tapeworm, whose prevalence was suppressed. Overall, our results provide a detailed case study illustrating the capacity for gene flow between allopatric populations of a given species to drive rapid adaptation, with an unusually fine-grained view of the earliest years of that introgression.

## Acknowledgments

We thank Jacqueline Salguero, Will Shim, Mariah Kenney, Joseph Marini for assistance with experiments (animal care, lab work). Discussion with Carl Veller greatly enhanced our understanding on how to estimate the effective strength of selection. Early versions of this work benefitted from constructive feedback from Catherine Peichel, Rowan Barrett, Dolph Schluter, Jesse Weber, Andrew Hendry, and Natalie Steinel.

## Author contributions

Conceptualization: DIB, FP, AP, BAF.

Methodology: DIB, FP, AP, BAF, SQ.

Investigation: FP, AR, AP, BAF, SQ, LS, MM, FG, AH, DIB

Visualization: BAF, DIB

Funding acquisition: DIB

Project administration: DIB, AR, BAF

Supervision: DIB

Writing – original draft: DIB, BAF, AP

Writing – review & editing: AP, BAF, FP, SQ, AR, FG, LS, MM, NR, AH, DIB.

## Data and materials availability

All data required to reproduce results of this paper have been deposited in a public database (sequence data have been deposited as two NCBI BioProjects (PRJNA1303581 and PRJNA1303714); other data and R scripts are publicly available at Zenodo (DOI: 10.5281/zenodo.16969467).

## Supplementary Materials

Materials and Methods

Figs. S1 to S7

Tables S1 to S4

**The PDF file includes:**

Materials and Methods

Figs. S1 to S7

Tables S1 to S4 References

## Materials and Methods

### Field collections

Wild caught threespine stickleback (*Gasterosteus aculeatus*) on Vancouver Island were collected using unbaited minnow traps set in the littoral zone during late May to early June across a 17-year period between 2005 to 2022 (years and sample sizes in Table S1). All collections were conducted under approval from the University of Texas at Austin IACUC (protocol AUP-03120501, AUP-05111701, AUP**-**07-032201 AUP-2010-00024, AUP-2015-00200), and subsequently the University of Connecticut IACUC (protocol A-18-008, A-21-025) and Scientific Fish Collection Permits from the Ministry of the Environment of British Columbia (NA05-19046, NA06-21423, NA07-32612, NA08-43012, NA09-34960, NA10-61026, NA11-70031, NA12-77018, NA13-85103, NA14-93580, NA15-16384, NA15-217759, NA18-287894, NA22-679623). Further details of the survey, including collection location coordinates are provided in (*18*, *24*, *25*, *38*). Upon capture, fish were euthanized in the field in MS-222. Fin clips were retained in ethanol, enabling subsequent genotyping work (see below). Specimens were either dissected when freshly caught (e.g., in 2022), or more typically were preserved in 10% neutral buffered formalin for subsequent dissection.

### Sample collection for DNA extraction and genotyping

Upon capture, fish were euthanized in the field, and a small portion of the caudal fin was clipped, placed in 95% ethanol, and stored at –20°C until DNA extraction. Fin clips used for whole genome sequencing were collected from fish sampled across the following lake-year combinations: Boot 2023, Comida 2009 and 2023, Gosling 2007, 2010, 2011, 2012, 2013, 2015, 2018, and 2022, Higgens 2009, Mohun 2009, Roberts 2013, and Sayward 2009. DNA was extracted using the Qiagen DNeasy Blood & Tissue Kit (catalog #69506), following the manufacturer’s instructions. For certain lab-raised individuals used in genotyping assays (e.g., validation of *spi1b* knockouts), DNA was extracted from freshly clipped fin tissue using the same protocol.

### High Coverage Sequencing of 2022 Gosling Lake Samples

We performed high-coverage whole genome sequencing (WGS) on 108 individual threespine stickleback collected from Gosling Lake in 2022. DNA was extracted from ethanol-preserved fin clips as described above, and libraries were prepared using the Illumina Nextera DNA Flex Library Prep Kit. Samples were pooled and sequenced in a single lane of an Illumina NovaSeq S4 v1.5 flow cell (300 cycles) at the Center for Genome Innovation (CGI), University of Connecticut, generating 150 bp paired-end reads. The sequencing design targeted an average depth of approximately 12–15× coverage per individual.

Reads were aligned to the *G. aculeatus* v.5 reference genome (*39*) using bwa -mem v.0.7.17 (Li, 2013). To tag PCR and optical duplicate reads we used the MarkDuplicates tool from the Picard package v.2.27.4. Variant calling was performed with GATK v.4.2.3 (DePristo et al., 2011), using HaplotypeCaller in GVCFmode for each individual separately, followed by joint genotyping using GenotypeGVCFs with the --include-non-variant-sites option. For variant filtering we generated a ‘callability mask’, identifying the genomic regions where we were unable to call variants with confidence. The callability mask consisted of four elements: (i) sites with unusually high or unusually low overall coverage, based on examining coverage histograms; (ii) sites where more than 10% of individuals had missing genotypes; (iii) sites identified by GATK as low quality (i.e., with the LowQual tag) and (iv) sites with ‘poor mappability’. Mappability was assessed by breaking down the genome into overlapping k-mers of 150bp (matching the read length) and mapping these k-mers back to the genome. Using the SNPable tool (http://lh3lh3.users.sourceforge.net/snpable.shtml), we then masked all sites where fewer than 90% of k-mers mapped back to their original location perfectly and uniquely. On autosomes, the callability mask comprised 94.8Mb, which is about 22.8% of sequence. After applying the callability mask, we added several hard filters based on GATK best practices: specifically focusing on mapping quality (MQ < 40), mapping strand bias (FS > 40), variant quality normalized by depth (QD < 2) and excess heterozygosity when compared with Hardy–Weinberg equilibrium (ExcessHet > 40).

To estimate the admixture proportions for the 108 Gosling Lake fish from 2022, we used PLINK v1.90b4.4 (*40*) with the resultant chromosome-specific VCF file as input and removed variable sites which had greater than 5% missing genotype call rate and only retained biallelic SNPs. Then genotype matrices were generated using PLINK v1.90b4.4 (*40*), and these genotypes were used to estimate allele frequencies and these data were combined with previously estimated allele frequencies from Gosling Lake sampled in 2009 (*24*) to calculate locus-specific Fst. For the top 5% most highly differentiated loci we ran a gene ontology enrichment analysis using the online platform g:Profiler (*41*) by, first, identifying the genes nearest the highly differentiated loci using bedtools (*42*). Then, because *Danio rerio* has more functional information than *G. aculeatus*, we identified orthologous genes in zebrafish and performed ontology enrichment analysis using the zebrafish orthologs. Then 2022 Gosling Lake genotypes were combined with previous ddRADSeq genotyping effort from Boot Lake in 2013 which is adjacent to Gosling Lake, and Comida Lake from 2013 (*43*). Then we performed a local PCA (*31*) to determine how population structure changes along the genome.

### Low Coverage Sequencing of Time Series Samples

We performed low-coverage whole genome sequencing (lcWGS; (*44*)) on 20 individuals per population from 15 lake-year combinations across Vancouver Island, including a time series from Gosling Lake spanning 2007 to 2018. Additional samples were collected from Boot, Comida, Higgens, Mohun, and Roberts Lakes, and Sayward Estuary; coordinates are provided in (*38*). Genomic DNA was extracted, (as mentioned above) and libraries were prepared using the Illumina Nextera DNA Flex Library Prep Kit, followed by sequencing on an Illumina NovaSeq S4 platform (paired-end 150 bp reads) at the Center for Genome Innovation (CGI), University of Connecticut. Sequencing targeted an average depth of ∼2X per individual.

Parsed paired-end reads assigned to individuals were mapped to the *G. aculeatus* reference genome (v.5 assembly; (*39*)) using BWA-MEM (*45*). Duplicate reads in the resultant alignments were identified and discarded using samblaster (*46*), and aligned reads were sorted, compressed, and indexed using samtools (*47*). Genotype likelihoods were computed in a probabilistic framework using ANGSD (*48*), implementing the GATK genotype likelihood model (-GL 2). Variable sites were identified via a likelihood ratio test, retaining sites with a significance threshold of *p* < 1e–6. We generated a genome-wide Beagle-format file, which was then split by chromosome for downstream analysis. To estimate ancestry and population structure, we applied PCAngsd (v1.35; (*49*)). Admixture proportions were calculated for each chromosome individually (K = 2), and a genome-wide covariance matrix was also computed from the same Beagle file, retaining the top four eigenvectors for principal components analysis.

PCAngsd estimated covariance matrix was used to perform a principal components analysis in R using the prcomp() function with a zero centered variable shift and scaled to unit variance. Then, to describe the genomic changes in Gosling Lake over time, we subset the prcomp() output to include Gosling Lake and Mohun watershed. Then for the Gosling Lake samples after the Mohun watershed fish introduction (> 2011), we calculated the mean PC1 and mean PC2 for each year. The Gosling Lake and Mohun watershed subset were then plotted as well as the year mean PC values with arrows connecting the estimated means. Then to more closely identify the introduction source, Admixture proportions (k = 3) were estimated for Gosling Lake fish from 2007 to 2018 and fish from the Mohun watershed including Comida Lake and Mohun Lake. We also estimated the covariance matrix for Mohun watershed popuoatio0ns and Gosling Lake fish from 2018, after the invasion. The subsequent covariance matrix was used to perfume a principal components analysis in R as described above and the first 40 PCs were used to perform a linear discriminate analysis in R using MASS::lda() with Mohun watershed populations (Comida Lake or Mohun Lake) as grouping variables. Then we predicted assignment to Comida or Mohun Lake for the 2018 Gosling Lake samples for all chromosomes individually. For the 2022 Gosling Lake samples the output .bed files were used as input for ADMIXTURE Version 1.3.0 (D.H. Alexander, J. Novembre, and K. Lange. Fast model-based estimation of ancestry in unrelated individuals. *Genome Research*, 19:1655–1664, 2009.) with K = 2. The PCAngsd and ADMIXTURE estimated admixture proportions for each chromosome were plotted using ggplot2::geom_density_ridges() for 2012, 2013 and 2022. Then for chr4 and chr7, the admixture proportions were plotted using ggplot2::geom_density_ridges() across years. The direction of Mohun watershed ancestry was inferred by including Comida Lake samples in the admixture proportion estimates.

We then plotted the trend in mean Mohun watershed ancestry for each chromosome across years. From this we also calculated a genome-wide average Mohun watershed ancestry for each year. This genome-wide trend was used to estimate the effective strength of selection acting on an average site in the genome. We fit a curve in which the between-year change in ancestry proportion should be equal to *s/2^g^*. Sampling a wide range of possible selection coefficients *s*, we calculated the sum of squares as a goodness of fit between the model prediction and the observed change in genome-wide mean ancestry. The best estimate of *s* is the value that minimized these sums of squares.

Within each sample year, we used an ANOVA to test whether the observed ancestry proportion differed among chromosomes. This allowed us to evaluate whether some chromosomes were, on average, introgressing faster or slower than others. Likewise, we used t-tests to evaluate whether a given chromosome increased (or, decreased) its ancestry proportion between years.

### Changes in *Spi1b^del^* frequency

Weber et al. (***24***) described a deletion in the gene *spi1b* that was associated with loss of fibrosis and increased tolerance of *S. solidus* infection in Gosling Lake stickleback. To assay changes in this focal gene over time, targeted genotyping to detect the *Spi1b^del^* deletion allele was performed using PCR with the following primers: forward 5′– tcactgagaaagcgccagtt–3′ and reverse 5′–tgtttcatggatgaatgaag–3′. The expected product size was 825 bp for wild-type alleles (*Spi1b+*) and 747 bp for alternate allele that had 78 bp intronic deletion (*Spi1b^del^*). PCR products were run on an agarose gel, stained with Cybersafe, and photographed. Comparison with a standard ladder allowed us to score individuals as wild type homozygote, deletion homozygote, or heterozygote. We used this PCR protocol to score *spi1b* genotype on individuals from 2007 through 2022 (sample sizes listed in Table S1). We used a binomial GLM to test for changes in allele frequency between years, and over the duration of the study (time as a numeric variable). The 2022 Gosling Lake called genotypes were combined with Comida Lake and Boot Lake genotyped using ddRADseq by Stuart et al (*18*). We then performed a principal components analysis as previously described with the fill of the points was based on *Spi1b* PCR genotypes.

### Functional validation of *Spi1b* role in fibrosis response: Spi1b knockout

*CRISPR/Cas9 Knockout of Spi1b* To disrupt *Spi1b*, we designed two sgRNAs targeting exon 2 using CHOPCHOP (*50*, *51*) based on the *G. aculeatus* BROADS1 genome assembly. The sgRNAs (5′–cgctcaccctcacgttcacc–3′ and 5′– acaggacgccatacggtacg–3′) were ordered as crRNAs from Integrated DNA Technologies (IDT). Each sgRNA was tested separately in independent microinjection experiments, and both produced comparable knockout efficiency.

RNP complexes were prepared by combining crRNA, tracrRNA, and Cas9 protein (all from IDT). Duplex buffer was used to make 100 μM stock solutions of individual crRNAs and tracrRNA. To generate crRNA:tracrRNA duplexes (50 μM), equal volumes of crRNA and tracrRNA were mixed and annealed in a thermal cycler: 95°C for 5 minutes, cooled to 25°C at 0.1°C/s, held at 25°C for 5 minutes, and then held at 4°C. The resulting 50 μM duplex was diluted to 25 μM for use in injection mixes and stored at –20°C. Injection cocktails (4 μL) were prepared fresh on the day of injection and included 0.4 μL 25 μM crRNA:tracrRNA duplex, 1.2 μL 50 μM Cas9 protein, 1.2 μL nuclease-free water, and 0.4 μL of 0.5% phenol red. Control cocktails (4 μL) contained 1.2 μL Cas9 protein, 2.4 μL water, and 0.4 μL phenol red, maintaining Cas9 concentration identical to that of treatment groups. Each mix was incubated at 37°C for 5 minutes before microinjection and kept at room temperature. Approximately 2 nL of the injection mix was injected into the yolk or cytoplasm of one-cell stage embryos using borosilicate capillaries.

Microinjections were performed following in vitro fertilization. Stickleback eggs were obtained from mature females and fertilized using sperm prepared by macerating male testes in Hank’s solution. Up to 100 eggs were combined with 50 μL of sperm solution to ensure fertilization. Embryos were kept covered to prevent drying during early development. Injection materials were prepared while embryos developed to the one-cell stage, typically within 20–25 minutes. Following injection, embryos were transferred to Petri dishes containing stickleback water (prepared from artificial seawater mix, 10% sodium bicarbonate, and deionized water). All procedures were performed at room temperature. The injected embryos were F1 hybrids generated from a cross between Gosling and Roberts Lake fish collected in 2018. These injected F1s, which were expected to be mosaic for CRISPR-induced mutations, were raised to maturity and crossed to produce F2 offspring. We screened F2 individuals for *Spi1b* knockouts using gel band shift PCR followed by Sanger sequencing. Despite extensive screening, we recovered only monoallelic knockouts; no biallelic knockouts were identified. This may reflect an essential role for *Spi1b* during development, as complete knockout could be lethal. This interpretation is consistent with prior reports of lethality in *Spi1* knockout mice (*52*). Additionally, we observed elevated mortality in mosaic crosses, further suggesting potential developmental constraints on complete gene loss. Genotyping was performed using the following primers: forward 5′–ggtgatttctgtcctgttttgga–3′ and reverse 5′–gtgcacgaaggtcatgaagc–3′.

*Testing Fibrosis Response in Spi1b KO Fish.* To test whether *Spi1b* influences the fibrosis response, we injected F2 monoallelic knockouts and non-knockout siblings with either saline (PBS) or alum. Intraperitoneal injections consisted of 20 μL total volume: either 20 μL PBS or 10 μL of 2% Alumax Phosphate (OZ Biosciences) mixed with 10 μL PBS. Alum is a widely used immune adjuvant that induces leukocyte recruitment and peritoneal fibrosis ((*53*); N. Steinel, pers. comm). For each genotype, 9 fish were injected with PBS and 9 with alum. Fish were sampled at 7 and 28 days post-injection. Fibrosis scores from days 7 and 28 were analyzed using linear models with fixed effects for treatment (alum vs. PBS), genotype (monoallelic Spi1b knockout vs. wild type) and dissection timepoint. Interaction terms were included to assess whether treatment and genotype effects varied across timepoints, and whether the genotypes exhibited different responses to treatment. AIC model selection was used to simplify the model to a subset of empirically justified terms.

*Scoring fibrosis:* Fibrosis was evaluated during dissection by assessing adhesion formation within the peritoneal cavity, following the protocol first established by Hund et al (*34*). In healthy stickleback, visceral organs move freely and are assigned a fibrosis score of 0. Fibrosis is characterized by adhesions between organs or between organs and the peritoneal wall. Severity was scored on an ordinal scale from 0 to 4, where 1 indicated mild adhesions limiting organ movement, 2 indicated adhesions between organs, 3 indicated adhesions between organs and the peritoneal wall, and 4 indicated severe adhesions making dissection of the cavity difficult. Previous assays confirmed that these ordinal scores are highly repeatable among independent observers (r > 0.9; (*54*)).

### Functional validation of *Spi1b* role in fibrosis response: pharmacological Spi1b inhibition

Stickleback fish from the Kenai River Flats (KRF) population (Alaska) initiate irreversible fibrosis within days of alum injection, and this fibrosis persists for up to one year. Lab-raised KRF adult fish were divided into six treatment groups, with seven individuals per group. Three groups received intraperitoneal injections of the *Spi1b* inhibitor DB1976 (*37*) at doses of 1 nM, 2 nM, or 4 nM, each co-injected with alum on day 0. A fourth group received alum co-injected with DMSO to serve as a vehicle control for the inhibitor treatments. A fifth group received two 10 μL injections of DMSO alone to control for injection number. All fish were assessed for fibrosis development on day 11 post-injection. Fibrosis was scored as described above. Fibrosis severity was analyzed using a linear model with DB1976 dose as a predictor among alum treated group. We also fit models including both alum treatment and DB1976 dose to assess their individual and combined effects on fibrosis.

### Testing whether alum-induced fibrosis contributes to *S. solidus* resistance

Lab raised stickleback from Gosling Lake (bred in 2018) were experimentally exposed to *S. solidus* as described in detail in (*24*, *55*). Briefly, live tapeworms were obtained from naturally infected Gosling Lake stickleback collected in the field and shipped to the University of Connecticut. Tapeworms were dissected from euthanized hosts, and size-matched pairs were placed in nylon biopsy bags suspended in breeding media in a dark, warm, shaking water bath. Eggs that passed through the mesh were collected from the bottom of the breeding jars and stored at 4°C. Eggs were hatched at 17°C under a 12-hour light cycle and resulting coracidia were used to infect *Macrocyclops albidus* copepods. Two weeks after exposure, individual copepods were screened under a dissecting microscope, and only those visibly infected were used for fish exposures.

A total of 118 Gosling Lake stickleback were distributed into two tanks and assigned to one of two treatment groups: fibrosis-induced (via alum injection) or control (PBS injection). Injections were administered one week prior to parasite exposure. Fish in the fibrosis group received an intraperitoneal injection of 10 μL 2% Alumax Phosphate (OZ Biosciences) mixed with 10 μL PBS. Control fish received 20 μL PBS. Fish were food-deprived for 24 hours before infection. On the day of exposure, water flow was paused and each fish was offered 10 infected copepods.

Fish were left to feed for 6–8 hours in static water, after which circulation was resumed. Copepod consumption was not monitored individually, but tanks were filtered after the feeding period to confirm uptake. All fish were dissected two months post-infection. During dissection, fibrosis was scored as described above. Parasite load (infection intensity) and tapeworm mass (total and individual) were recorded. In cases where tapeworm mass fell below the limit of scale detection (<0.0001 g), individuals were assigned a mass of 0.00005g. Linear models were used to test whether total tapeworm mass depended on fibrosis severity. A Poisson generalized linear model (GLM) was used to test whether infection intensity depended on fibrosis score.

### Changing infection prevalence and fibrosis through time

Freshy caught or archived formalin-preserved stickleback were dissected to count *S. solidus* parasites. Prior to dissection we measured individuals’ standard length and body mass. During dissection we also recorded sex based on gonad anatomy. We include previously published data on *S. solidus* prevalence (*24*), supplemented with additional years of samples from archived stickleback specimens, and more recent fresh caught samples. To test for a directional temporal trend in infection prevalence, we fit a binomial generalized linear model (GLM) with year as a continuous predictor of the number of infected individuals out of the observed sample per year. We also tested for a correlation between tapeworm prevalence and *Spi1b^del^*allele frequency.

We dissected and scored fibrosis from archived formalin-preserved samples of stickleback originally collected 2005 and 2014, and fresh-caught samples collected in 2016 and 2022. We used the ordinal scale to score fibrosis as described above. Infection intensity was also noted for each individual. We used linear and quadratic regression to test whether mean fibrosis score exhibited a directional or curvilinear trend through time. We then tested whether mean fibrosis was related to *S. solidus* prevalence, treating each year as an observation and considering both linear and quadratic relationships. *A priori* we considered a quadratic relationship likely based on a recent theoretical model (*56*). Within each year, we used a correlation test to evaluate whether fibrosis severity was related to infection intensity. Our expectation was that before introgression, fibrosis should be mostly absent and unrelated to infection (consistent with prior work by Weber et al (*24*). However, once the immigrant *Spi1b^+^*genotype invaded we expected to find a positive relationship between infection and fibrosis, consistent with laboratory evidence that infection induces fibrosis.

**Figure S1:**
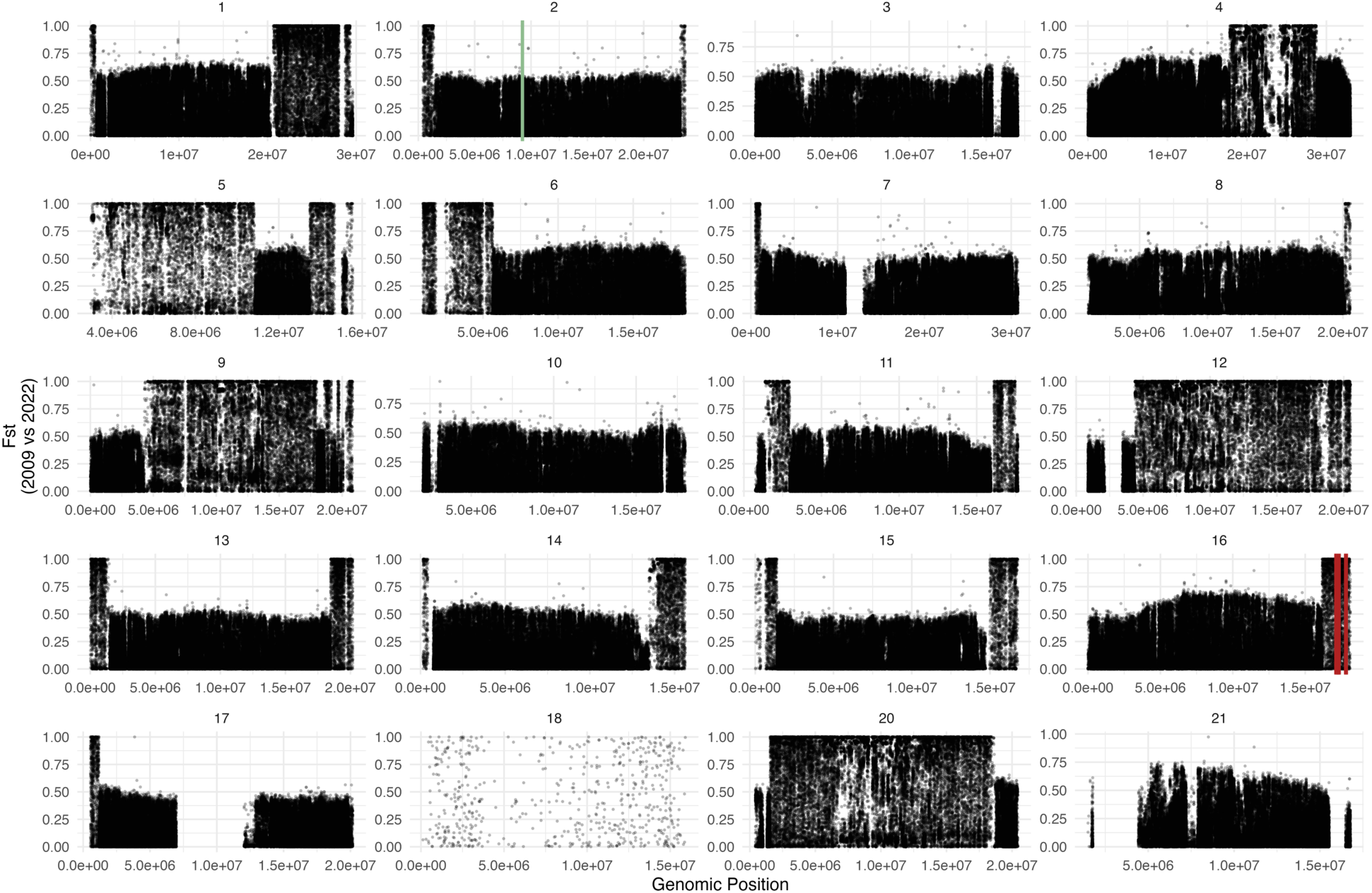
Genome-wide allele frequency change between 2009 and 2022 Gosling Lake stickleback. Genetic divergence in allele frequencies (FST) between Gosling Lake stickleback sampled in 2009 (N = 100 individuals genotyped by PoolSeq to obtain allele frequencies at an average of 2x depth), versus Gosling Lake stickleback sampled in 2022 (N = 108 whole genome sequences, at an average of 10x depth). Each panel is a chromosome, omitting the Y and X chromosomes, with Spi1b genomic position indicated in green on Chromosome 2 and Trace-amine associated receptor loci in red on Chromosome 16.

**Figure S2:**
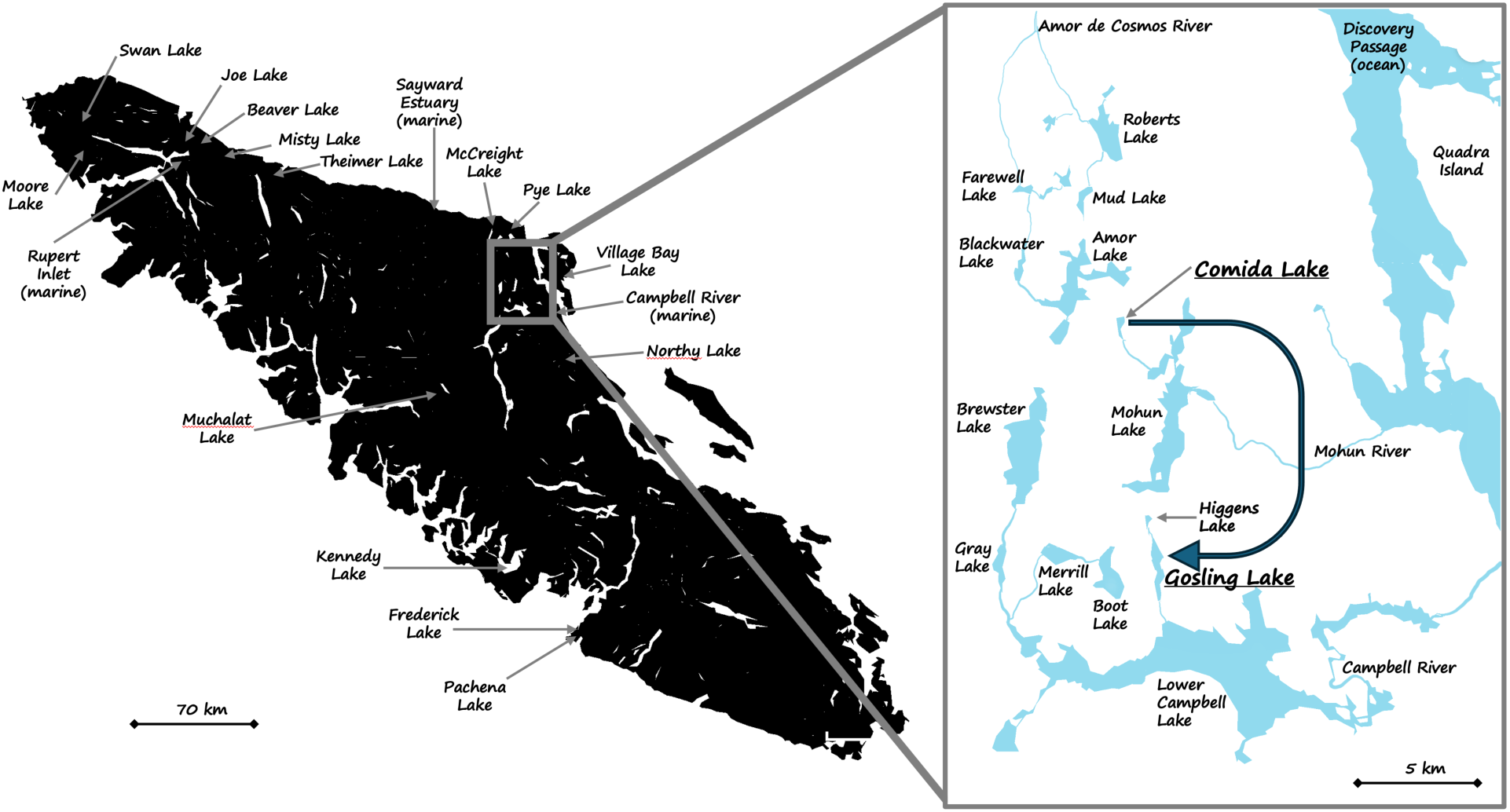
A map of Vancouver Island showing the populations examined here with a detailed inset. For the larger map of the island, we provide names and arrows pointing to the populations genotyped using ddRADseq by Stuart et al (*18*), which we use to evaluate possible sources of introgression. A complete phylogeny of these populations can be seen in (*18*). In the inset we show a subset of the lakes in the region, which are divided into three major watersheds: Amor de Cosmos River draining north, Mohun River, and Campbell River. Gosling Lake, the focal population in this study, is indicated with larger font, and is in the Campbell River drainage. Comida Lake, the source of migrants (also larger font), is in the Mohun River drainage. Roberts Lake (top) is in the Amor de Cosmos drainage and was used in previous crosses with Gosling Lake fish that identified *spi1b* as the likely genetic basis of variation in fibrosis based immunity to *S. solidus* infections.

**Figure S3:**
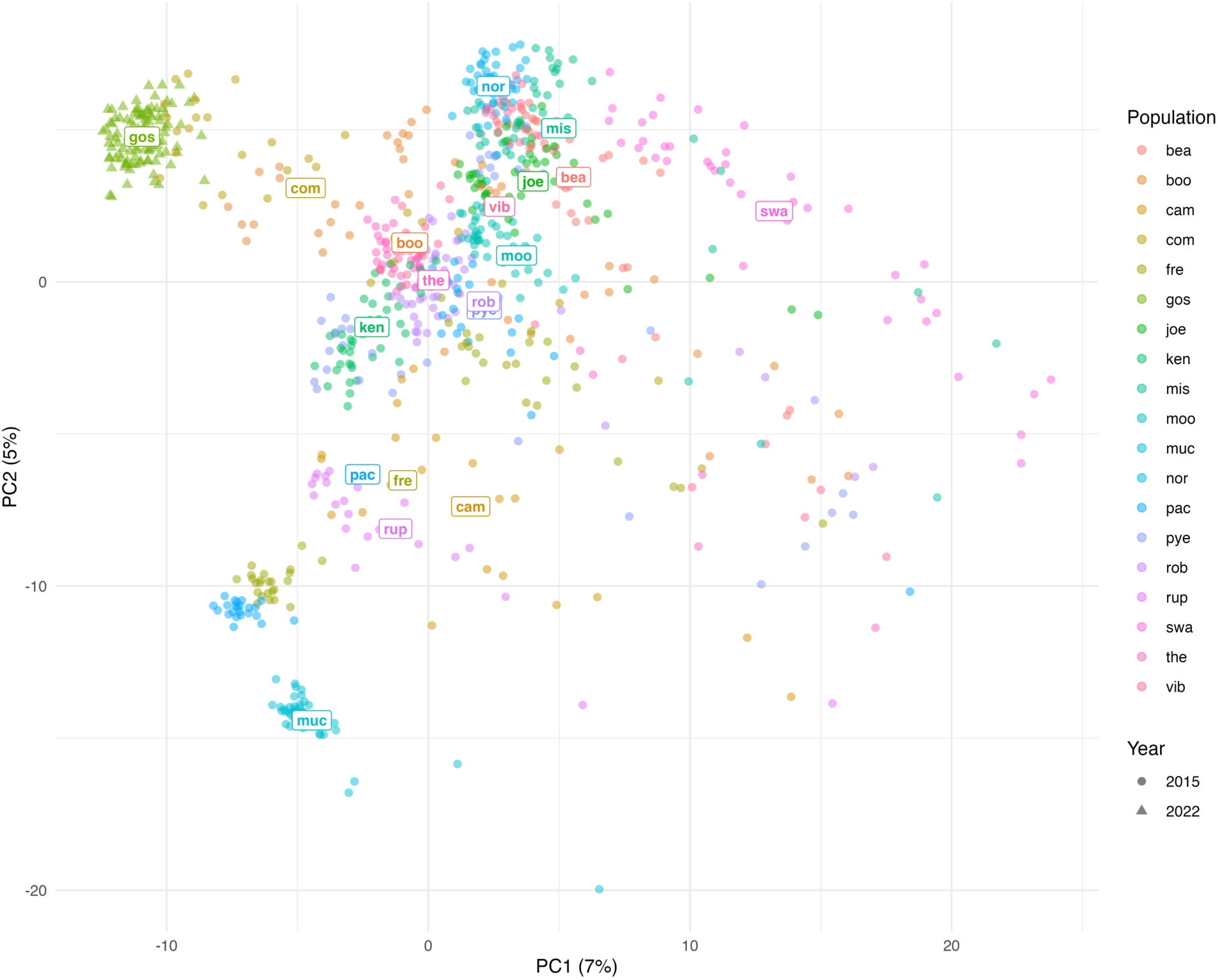
Principal component analysis of population differentiation using 2022 genomic SNPs and 2015 ddRAD data. Population differentiation PCA using 2022 genomic sequence SNPs and ddRAD data from 2015 (from the study by Stuart et al, (*43*)). All populations shown here are represented in the map in Fig. S2: bea = Beaver Lake, boo = Boot Lake, cam = Campbell River Estuary marine fish, com = Comida Lake, fre = Frederick Lake, gos = Gosling Lake, joe = Joe Lake, ken = Kennedy Lake, mis = Misty Lake, moo = Moore Lake, muc = Muchalat Lake, nor = Northy Lake, pac = Pachena Lake, pye = Pye Lake, rob = Roberts Lake, rup = Rupert Inlet marine fish, swa = Swan Lake, the = Theimer Lake, vib = Village Bay Lake. GPS coordinates for all populations are listed in data supplements in (*18*).

**Figure S4:**
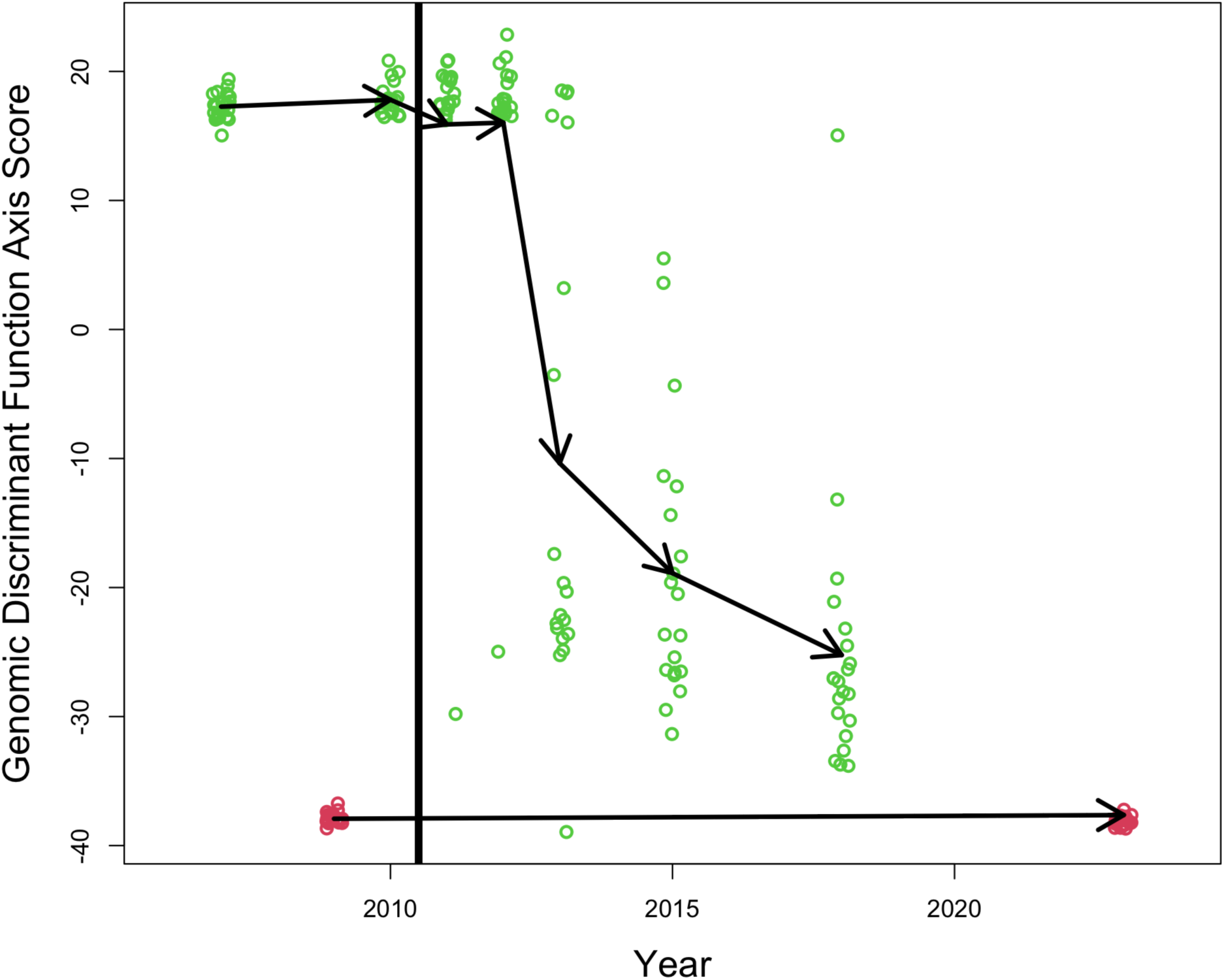
Discriminant function analysis reveals Comida Lake introgression into Gosling Lake over time. Alternative visualization of introgression. The red points are Comida Lake individuals, and green points are Gosling Lake individuals. We trained a discriminant function analysis based on pre-2011 samples to distinguish the two lakes, then applied this function to post-2011 samples to predict their ancestry contributions. This analysis revealed one individual fish sampled in 2011 and one in 2012 with Comida Lake ancestry contributions. In 2013, there is a mix of pure Gosling and hybrid individuals. But by 2015 no pure Gosling genotypes were sampled (possibly one sampled in 2018), revealing the systemic spread of the foreign Comida Lake genotypes.

**Figure S5:**
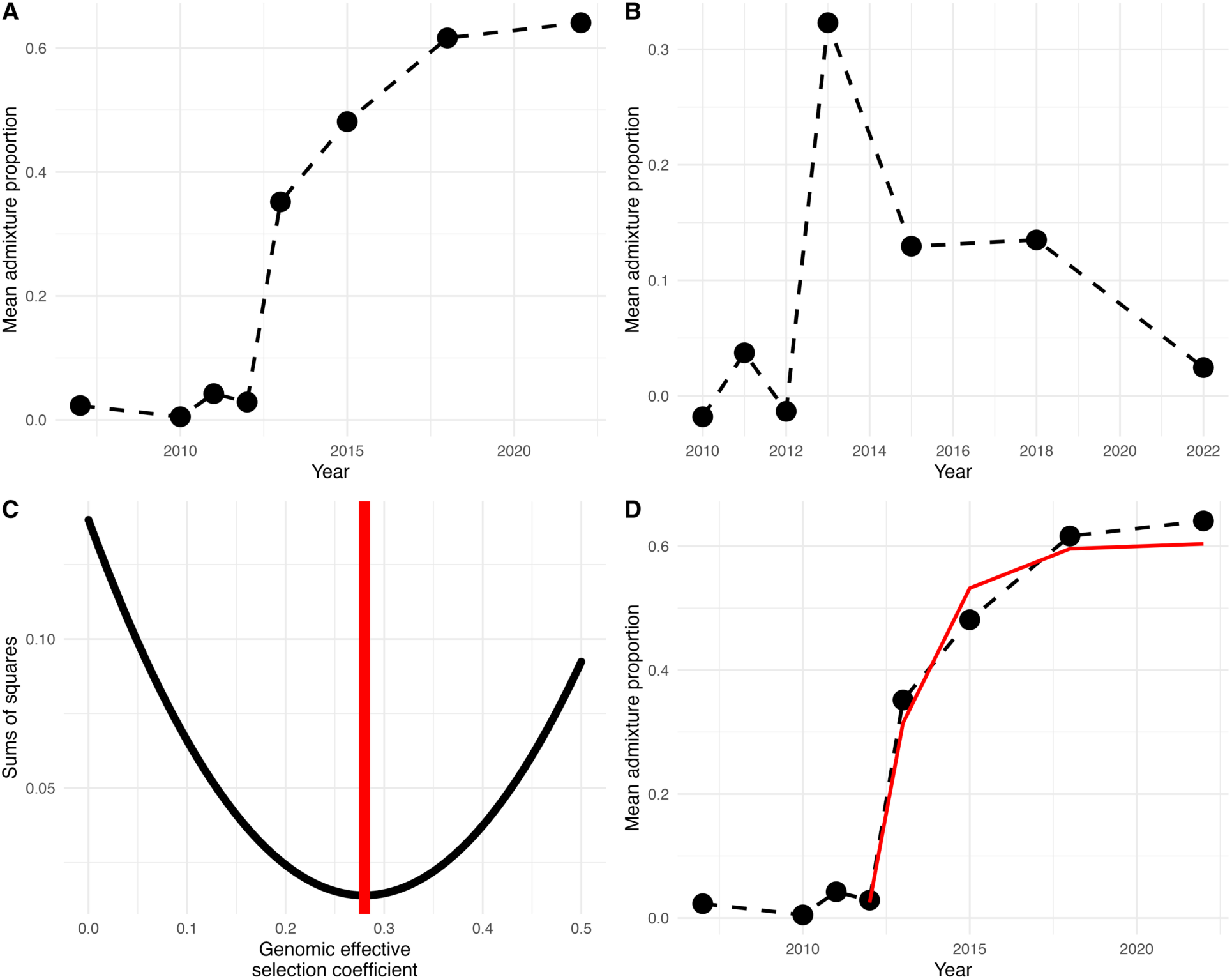
Estimating genome-wide effective selection coefficients. A) Observed data on the estimated admixture proportion (% Mohun watershed ancestry), averaged across the genome and across individuals. B) Observed change in admixture proportion between time points. C) We fit a model with the change in admixture proportion between successive years equal to s/2^g^. That is, in the first generation the entire genome experiences selection coefficient *s*. In the next generation, s/2, then s/4, then s/8. Where we missed a year of sampling, the observed change in admixture proportion would be the sum of the expected values. We then calculated the sums of squares as a measure of goodness of fit between our model, and the observed values in (B), for a variety of values of *s*. We plot a vertical red like at the estimate of *s* that minimizes the sums of squares (maximum fit to the data). (D) We then replot the data from panel (A) with the modelled allele frequency change for a neutral locus experiencing hitchhiking overlain in red.

**Figure S6:**
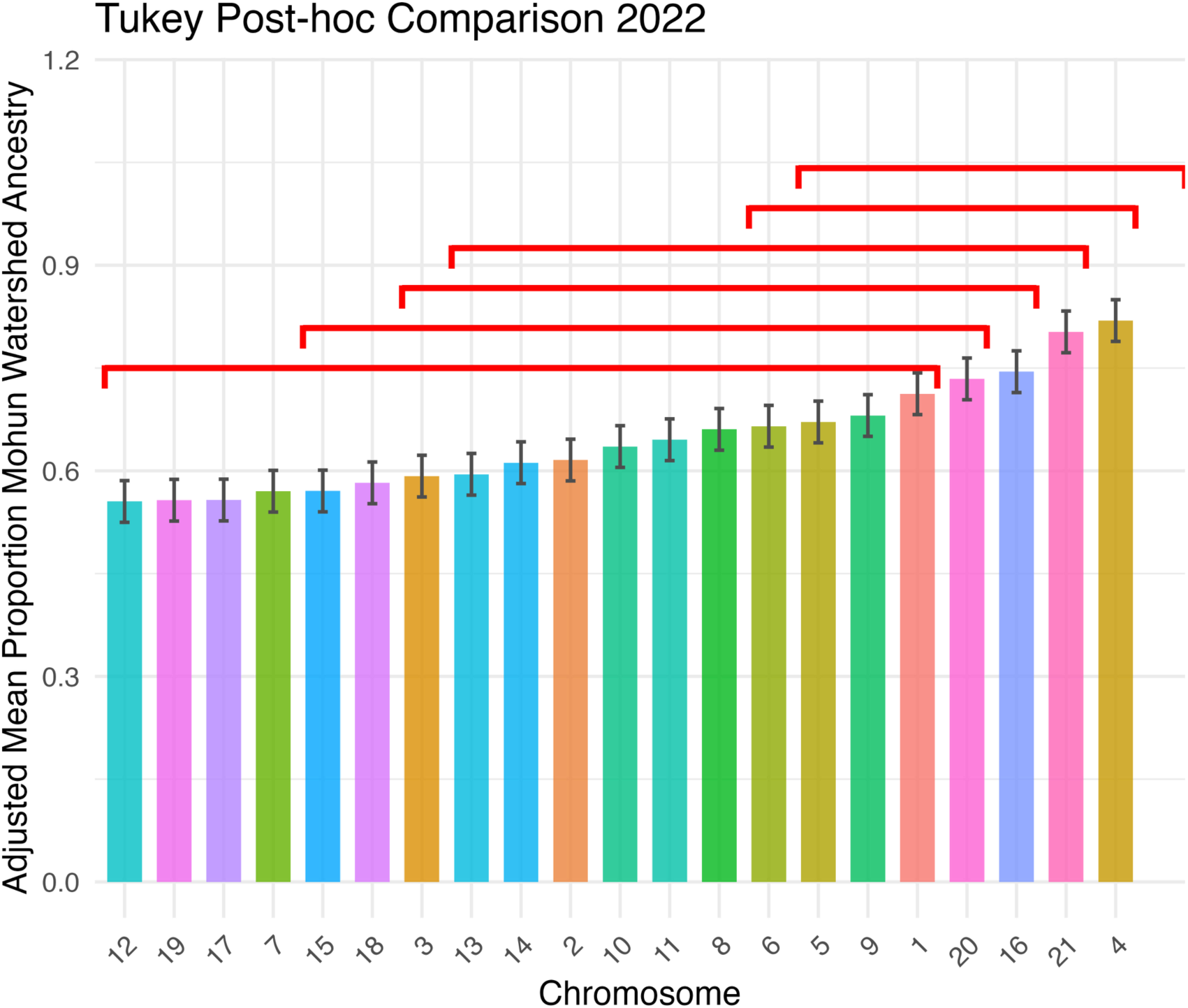
Chromosome-specific differences in Mohun watershed ancestry in 2022 Gosling Lake stickleback. Among-chromosome differences in ancestry proportion in the 2022 sample from Gosling Lake, using an ANOVA to test for overall differences, and Tukey Post-Hoc tests for pairwise differences the red brackets indicate group membership for post-hoc tests. Equivalent tests from 2005-2018 showed no significant differences in ancestry between chromosomes (all P>0.05). The relatively low introgression on the Y chromosome and X chromosome (Chr19) suggest that sexual selection (e.g., inherent fecundity or mate attraction differences) are not driving the introgression documented here.

**Figure S7:**
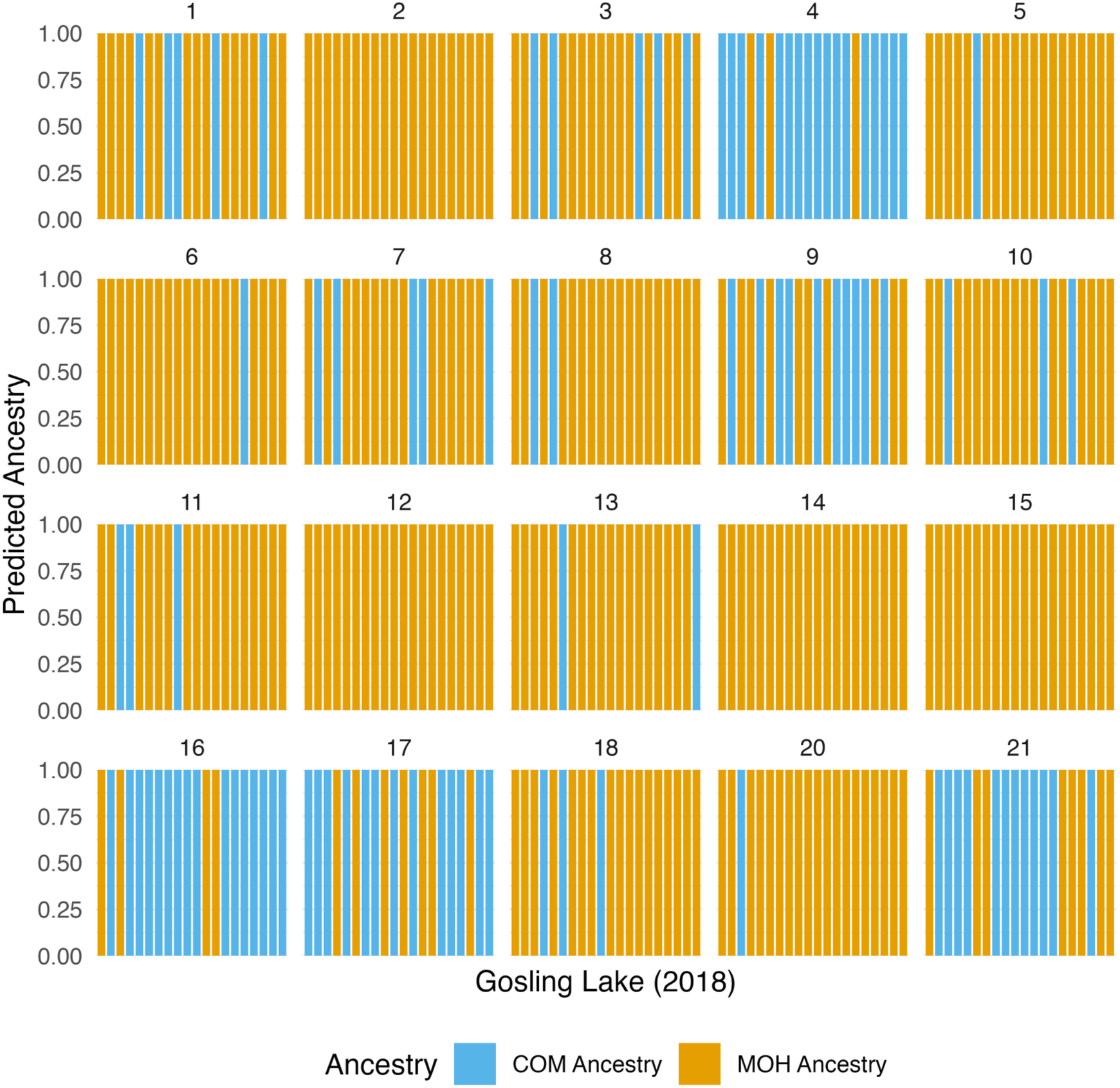
Linear discriminant analysis of principal components. The predicted population membership of post-invasion Gosling Lake individuals sampled in 2018 to the potential invasion sources from the Mohun watershed (COM/MOH) across all chromosomes. The LDA was trained on a PCA with the hypothesized invasion sources as the grouping variables then used to predict 2018 Gosling Lake individual invasion source membership.

**Table S1:** Sample sizes by lake and year for low coverage whole-genome sequencing. Lake-year combinations and the number of individual stickleback sampled for whole-genome sequencing at approximately 2X coverage. The 108 individuals sampled from Gosling Lake in 2022 (not shown here) were sequenced at higher depth (∼10X coverage).

| **Sampling Year** | **Population** | **Sample Size for low coverage WGS** |
| --- | --- | --- |
| 2023 | Boot Lake | 19 |
| 2023 | Comida Lake | 20 |
| 2009 | Comida Lake | 19 |
| 2007 | Gosling Lake | 21 |
| 2010 | Gosling Lake | 20 |
| 2011 | Gosling Lake | 20 |
| 2012 | Gosling Lake | 20 |
| 2013 | Gosling Lake | 19 |
| 2015 | Gosling Lake | 20 |
| 2018 | Gosling Lake | 21 |
| 2022 | Gosling Lake | 108* |
| 2009 | Higgins Lake | 22 |
| 2009 | Mohun Lake | 23 |
| 2013 | Roberts Lake | 22 |
| 2009 | Sayward Estuary | 23 |

\* Used for 10x coverage whole genome sequencing

**Tables S2:**
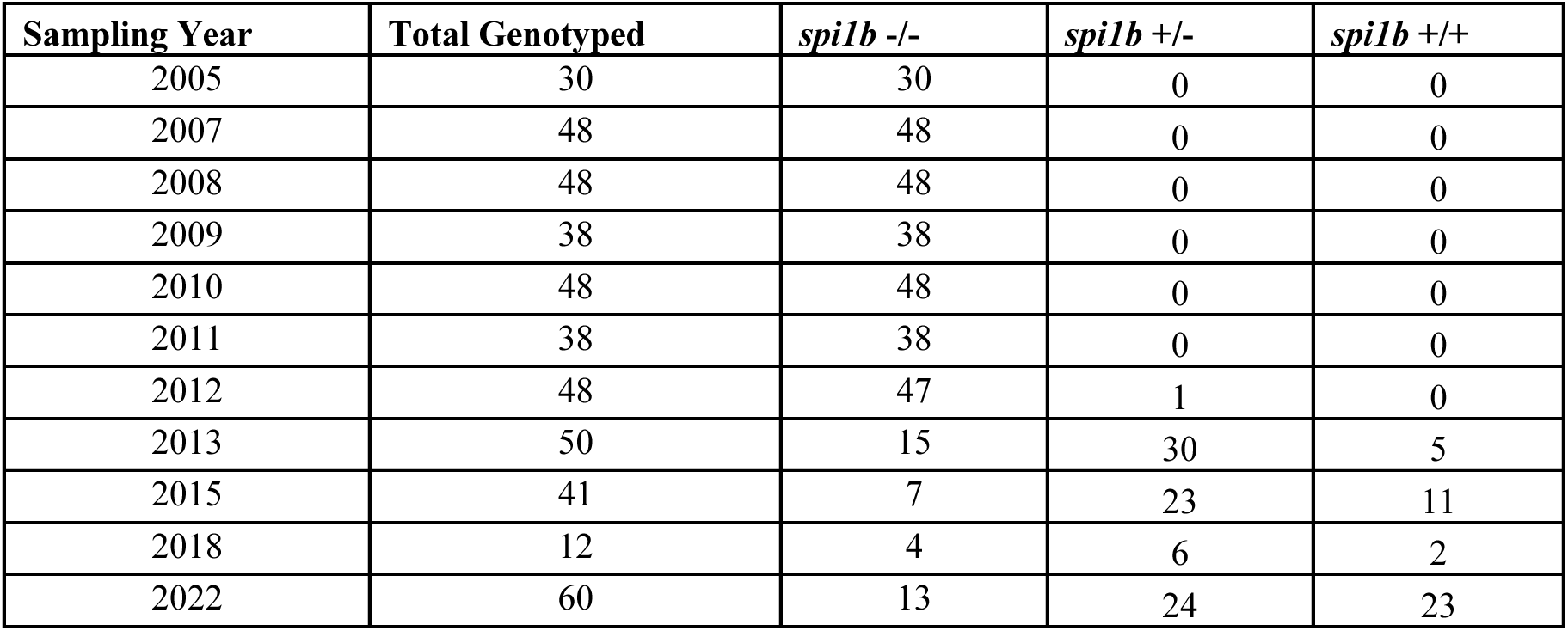
Genotype counts at the spi1b locus in Gosling Lake from 2005 to 2022. Total number of individuals from Gosling Lake genotyped for the *spi1b* locus by PCR from 2005 to 2022. Genotypes are categorized as *spi1b* -/- (homozygous for the deletion), *spi1b* +/- (heterozygous), and *spi1b* +/+ (homozygous for the full-length allele). PCR genotyping was performed on fin clips using a custom primer set, as described in the Methods.

**Table S3:** Prevalence of S. solidus infections in Gosling Lake stickleback across years. Sample sizes and number of S. solidus infected individuals dissected from Gosling Lake stickleback across multiple years. Years marked with * indicate samples that were dissected from archived formalin preserved fish from Gosling Lake.

| **Year** | **N dissected** | **N infected** |
| --- | --- | --- |
| 2005* | 33 | 14 |
| 2006 | 435 | 318 |
| 2009 | 147 | 81 |
| 2010 | 21 | 10 |
| 2012 | 17 | 10 |
| 2013* | 39 | 15 |
| 2014 | 84 | 13 |
| 2015 | 60 | 35 |
| 2016 | 31 | 13 |
| 2022 | 145 | 18 |

**Table S4:** Frequency of fibrosis in dissected Gosling Lake stickleback across years. Number of Gosling Lake stickleback dissected and recorded as fibrotic across sampling years. Years marked with * indicate samples that were dissected from archived formalin preserved fish from Gosling Lake.

| Years | N dissected | N fibrotic |
| --- | --- | --- |
| 2005* | 33 | 2 |
| 2013* | 39 | 20 |
| 2014* | 33 | 11 |
| 2022 | 145 | 6 |

